# Common genetic variants contribute to risk of rare severe neurodevelopmental disorders

**DOI:** 10.1101/309070

**Authors:** Mari E. K. Niemi, Hilary C. Martin, Daniel L. Rice, Giuseppe Gallone, Scott Gordon, Martin Kelemen, Kerrie McAloney, Jeremy McRae, Elizabeth J. Radford, Sui Yu, Jozef Gecz, Nicholas G. Martin, Caroline F. Wright, David R. Fitzpatrick, Helen V. Firth, Matthew E. Hurles, Jeffrey C. Barrett

## Abstract

There are thousands of rare human disorders caused by a single deleterious, protein-coding genetic variant ^1^. However, patients with the same genetic defect can have different clinical presentation ^2–4^, and some individuals carrying known disease-causing variants can appear unaffected ^5^. What explains these differences? Here, we show in a cohort of 6,987 children with heterogeneous severe neurodevelopmental disorders expected to be almost entirely monogenic that 7.7% of variance in risk is attributable to inherited common genetic variation. We replicated this genome wide common variant burden by showing that it is over-transmitted from parents to children in an independent sample of 728 trios from the same cohort. Our common variant signal is significantly positively correlated with genetic predisposition to fewer years of schooling, decreased intelligence, and risk of schizophrenia. We found that common variant risk was not significantly different between individuals with and without a known protein-coding diagnostic variant, suggesting that common variant risk is not confined to patients without a monogenic diagnosis. In addition, previously published common variant scores for autism, height, birth weight, and intracranial volume were all correlated with those traits within our cohort, suggesting that phenotypic expression in individuals with monogenic disorders is affected by the same variants as the general population. Our results demonstrate that common genetic variation affects both overall risk and clinical presentation in disorders typically considered to be monogenic.

---

We carried out a genome-wide association study (GWAS) in 6,987 patients with severe neurodevelopmental disorders and 9,270 ancestry-matched controls, using common variants with a minor allele frequency ≥5% (Figure 1, Extended Data Figure 1, Supplementary Tables 1-2 and Methods). The patients were recruited in the UK and Ireland as part of the Deciphering Developmental Disorders (DDD) study ^6,7^, by senior clinical geneticists who had assessed their developmental disorder was of sufficient severity that it was likely monogenic. In addition to neurodevelopmental defects (*e.g*. global developmental delay, intellectual disability, cognitive impairment or learning disabilities in 86%, autism spectrum disorders in 16%, Figure 2a), 88% also had abnormalities in at least one other organ system (Figure 2b and Extended Data Table 1).

**Figure 1.**
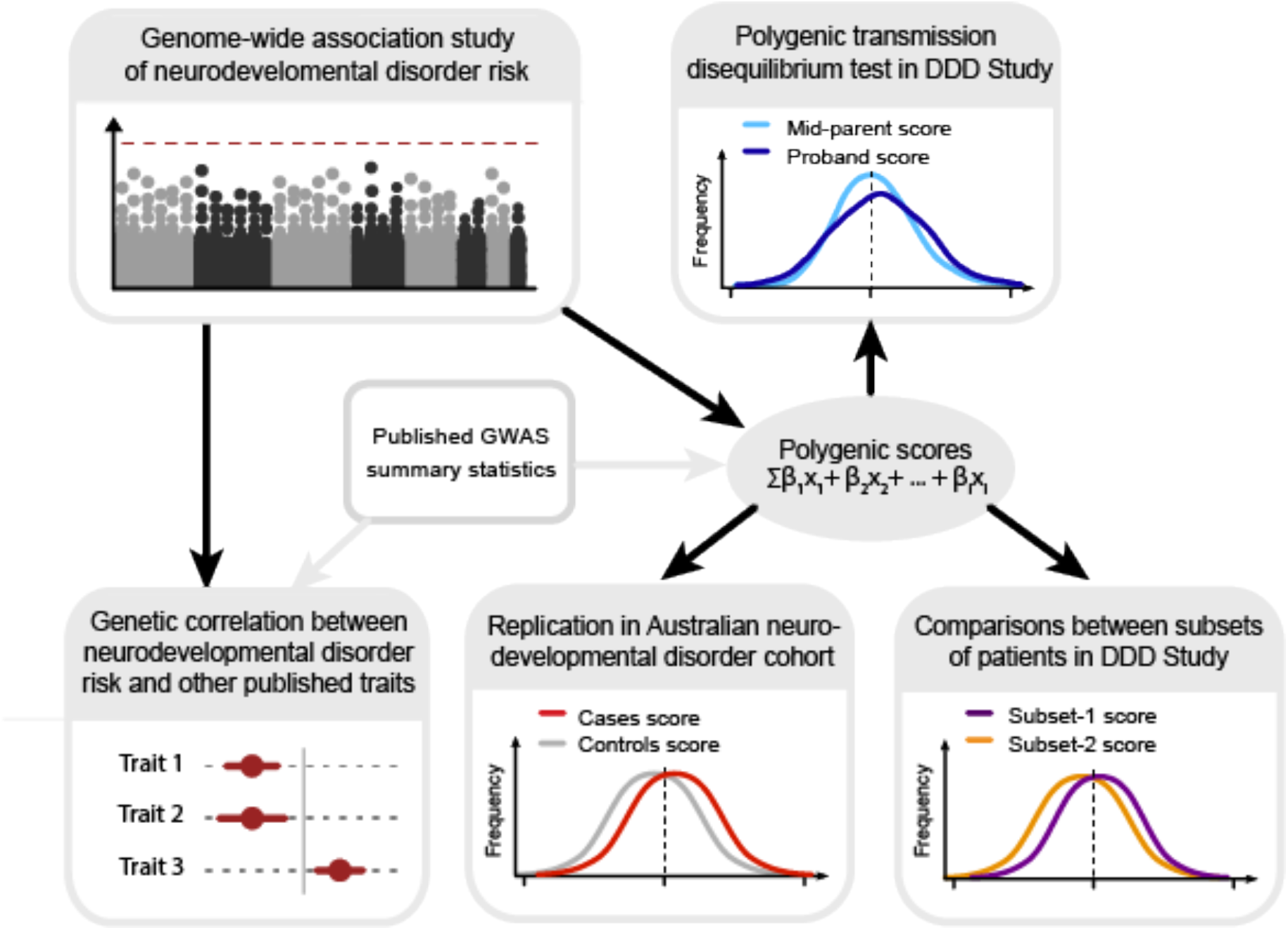
Outline of our analysis exploring the contribution of common variants to risk of severe developmental disorders. We do this by first conducting a discovery GWAS in a large dataset of neurodevelopmental disorder patients, and we validate our findings through analysis of polygenic transmission in independent trios from the same cohort. We then investigate how polygenic risk for neurodevelopmental disorders compares to published studies in neuropsychiatric and other traits via genetic correlation, and further replication in an independent Australian cohort. Finally, we explore how polygenic effects are distributed in our cohort and their contribution to specific phenotypes.

**Figure 2.**
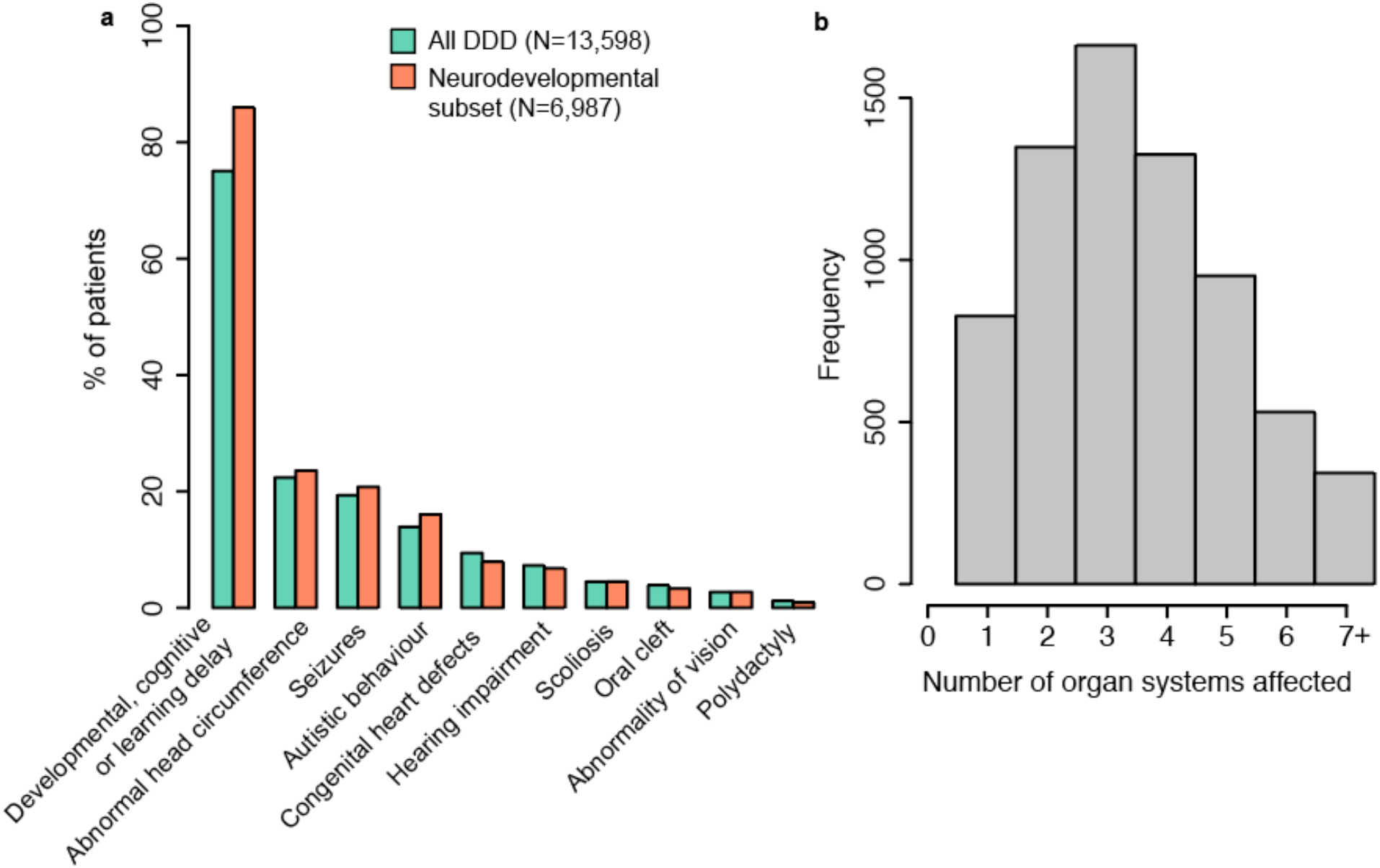
Patients recruited to the DDD study have diverse phenotypes. A. Examples of specific phenotypes affecting different organ systems, observed in the full DDD cohort and the neurodevelopmental subset of patients. B. Distribution of the number of distinct organ systems affected in the set of 6,987 patients with neurodevelopmental abnormalities (Methods).

We did not find any single variant associations at genome-wide significance (Extended Data Figure 2a), which was unsurprising given the heterogeneity of our clinical phenotype and the presumption that these disorders are monogenic. We did, however, observe a modest inflation in the test statistics (λ=1.097, Extended Data Figure 2b), which could indicate either residual bias between cases and controls, or evidence of a polygenic contribution of common variants to disease risk. We therefore estimated common variant heritability using LD score regression^8^, which can differentiate between these two possibilities, and found that 7.7% (SE=2.1%) of variance in risk (on the liability scale) for neurodevelopmental disorders in our sample was attributable to common genetic variants, when assuming a population prevalence of 1% (Methods). This common variant heritability estimate (h^2^) is similar to what has been reported for common disorders such as autism (h^2^=11.8%, SE=1.0%) ^9^ and major depressive disorder (h^2^=8.9%, SE=0.4%) ^10^. To replicate this signal, we analysed an independent set of 728 parent-child trios recruited as part of the same study, but who were not in the initial GWAS. We calculated polygenic scores for each individual by summing the genetic effects across all independent variants from our discovery GWAS (Figure 1 and Methods). We then performed a polygenic transmission disequilibrium test^11^, which compares the mean parental polygenic scores to those of the affected children. We found that our neurodevelopmental disorder risk score was overtransmitted in these trios (P=0.0035, t=2.48, df=727, one-sided t-test), confirming that common variants contribute to risk of disorders widely presumed to be monogenic.

Previous studies have shown that risk of more common neuropsychiatric disorders (e.g. schizophrenia and bipolar disorder ^12,13^) and variation in other brain-related traits, including educational attainment, ^13,14^ is driven in part by shared common genetic effects. We therefore used the LD score method ^15^ to test for genetic correlation between our neurodevelopmental disorder GWAS and available GWAS data for common neuropsychiatric disorders, cognitive and educational traits, anthropometric traits, and negative control diseases that have well powered GWAS but are not related to neurodevelopment. We found that genetic risk for neurodevelopmental disorders was significantly negatively correlated with genetic predisposition to higher educational attainment (r_g_=-0.49, SE=0.079, P=5.1x10”^10^) ^16^ and intelligence ^17^ (as measured by Spearman’s *g)* (r_g_=-0.44, SE=0.104, P=2.2x10^−5^), and positively correlated with genetic risk of schizophrenia (r_g_=0.29, SE=0.071, P=5.9x10^−5^) (Figure 3 and Extended Data Table 2). None of the anthropometric traits, nor the negative control traits, were significantly genetically correlated with our data, after accounting for multiple testing. We also used partitioned LD score regression ^18^ to show that heritability of neurodevelopmental disorders was nominally significantly enriched in cells of the central nervous system (P=0.02) (Extended Data Table 3), and in mammalian constrained regions ^19^ (P=0.009) (Supplementary Table 3), consistent with similar analyses for other neuropsychiatric and cognitive traits. Together, these results suggest that thousands of common variants have individually small effects on brain development or function, which in turn influences neuropsychiatric disease risk, cognitive traits, and risk for severe neurodevelopmental disorders.

**Figure 3.**
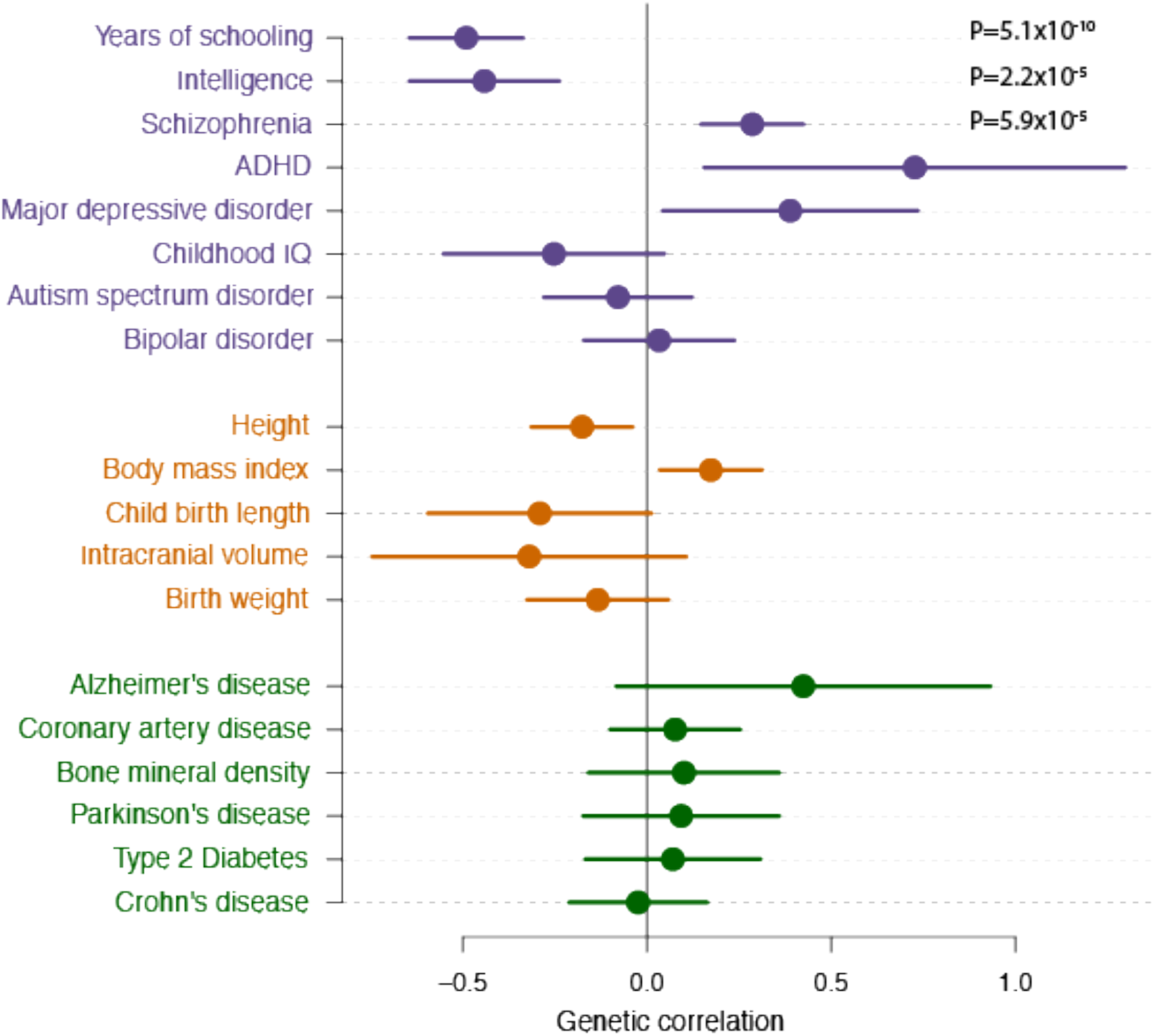
Genetic correlations between neurodevelopmental disorder risk and a range of traits. Cognitive/psychiatric traits (purple), anthropometric traits (orange) and negative control traits (green). Genetic correlation is calculated using bivariate LD score correlation ^15^, with the bars representing 95% confidence intervals (standard error) before correction for multiple testing. Traits annotated with a P-value pass Bonferroni correction for 19 traits.

We next investigated how general our genetic correlation findings were, by attempting to replicate them in another neurodevelopmental disorder cohort (Figure 1). We obtained GWAS data for 1,270 developmental disorder cases from Australia and 1,688 ancestry-matched Australian controls. Because this sample is too small to do direct genetic discovery, we tested common variant polygenic scores using summary statistics from our discovery GWAS and the published GWAS, including educational attainment ^16^ and intelligence ^17^. We replicated our observation of lower genetic scores for educational attainment and intelligence in the neurodevelopmental disorder cases compared to controls (P=1.4x10^−8^ and P=7.6x10^−4^ respectively), and found that cases had a nominally significantly increased score for schizophrenia (P=0.014) (Methods and Extended Data Table 4). We did not see a significant difference between cases and controls for the score constructed from our own discovery GWAS, which may be because our GWAS is relatively underpowered, due in part to much smaller sample sizes, compared to the GWAS studies of cognition, educational attainment and schizophrenia.

These findings could mean that common variants entirely explain a subset of patients with neurodevelopmental disorders, and are not relevant in the remainder, or that all patients’ disorders have both rare and common variant contributions (Figure 1). We have exome sequenced our cohort of patients, as well as their parents, and have previously reported a variety of both *de novo* and inherited diagnostic variants ^20,21^. We therefore compared polygenic scores for cognitive traits and neuropsychiatric disorders between patients for whom we had identified diagnostic or probably diagnostic variants in a known developmental disorder gene ^22^ (N=1,127) and those who had no candidate diagnostic variant (N=2,479), but we found no significant differences for any polygenic score we tested after controlling for multiple testing (Extended Data Table 5 and Methods). We showed by simulations that if the “diagnosed” cases had the same distribution of the educational attainment polygenic score as controls we would have had sufficient power to detect a difference between them and the undiagnosed cases (Methods). This is consistent with a previous study in autism ^11^ that similarly found no evidence for a difference in polygenic risk scores between autism cases with a *de novo* diagnostic mutation compared to those without. This suggests that both common and rare variants are contributing in many neurodevelopmental disorder patients. However, as the DDD project continues to identify new diagnoses, we anticipate that the increase in power may show that monogenic and polygenic contributions are not purely additive.

In addition to showing that common variation affects overall risk of severe neurodevelopmental disorders, we sought to determine if it can also affect individual presentation of symptoms. We identified four phenotypes measured in our neurodevelopmental disorder cohort for which independent GWAS data are available: autism (16% of cohort), birth weight, height, and intracranial volume. On average, our neurodevelopmental patients had a head circumference 1.20 standard deviations (SD) smaller, they were 0.72 SD shorter than, and weighed 0.15 SD less than the age and sex-adjusted population average. We constructed common variant polygenic scores for the four phenotypes as described above, and tested for association between the relevant score and phenotype in our cohort. In all four cases, there was significant association (Table 1 and Extended Data Table 6), demonstrating that common variation contributes to the expression of these traits in our study. We next tested for association between the educational attainment polygenic score and severity of overall neurodevelopmental phenotype. We found that patients with severe intellectual disability or developmental delay (N=911, Methods) had higher scores (i.e. greater educational attainment, proxy for higher cognitive function, P=0.003, Table 1) than those with mild or moderate disability or delay (N=1,902). This finding, which might seem initially counter-intuitive, is consistent with epidemiological studies ^23^ which found that the siblings of patients with severe intellectual disability showed a normal distribution of IQ, whereas siblings of patients with milder intellectual disability had lower IQ than average, implying that mild intellectual disability represents the tail-end of the distribution of polygenic effects on intelligence and severe intellectual disability has a different etiology.

**Table 1.** Polygenic score analyses in the DDD Study.

| Measured trait | Polygenic score | Results* |  |  |  |
| --- | --- | --- | --- | --- | --- |
|  |  | Beta | Standard error | P-value | R<sup>2</sup> |
| Birth weight (N=6,496) | Birth weight | 0.187 | 0.017 | 2.55x10<sup>-28</sup> | 0.020 |
| Height (N=5,465) | Height | 0.408 | 0.033 | 1.18x10<sup>-35</sup> | 0.033 |
| Head circumference (N=6,074) | Intracranial volume | 0.132 | 0.031 | 1.79x10<sup>-5</sup> | 0.004 |
| Autistic behavior: affected (N=1,121), unaffected (N=5,866) | Autism spectrum disorder | 0.120 | 0.033 | 2.53x10<sup>-4</sup> | 0.006 ‡ |
| Developmental delay or intellectual disability: severe (N=911), mild/moderate (N=1,902) † | Educational attainment | 0.117 | 0.040 | 0.003 | 0.009 ‡ |

\*Linear or logistic regression of measured traits in the DDD Study against the respective polygenic score, including ten ancestry principal components as covariates. † Severe cases were labelled as 1 in the logistic regression. ‡ Nagelkerke R<sup>2</sup>

The study of human disease genetics has often been segregated into rare, single gene disorders, and common complex disorders. There is abundant evidence that rare variants in individual genes can cause phenotypes seen much more commonly in individuals without a monogenic cause, including genes for maturity onset diabetes of the young ^24^, familial Parkinson’s disease ^25^, and breast cancer in carriers of *BRCA1* or *BRCA2* mutations. Here we have shown that the same interplay between rare and common variation exists even in severe neurodevelopmental disorders typically presumed to be monogenic. Previous studies have shown that the penetrance and expression of these disorders are affected by which specific missense variant is carried ^26^ and the presence of mutations in secondary modifier genes ^27^. Here, we have demonstrated that they are also modified by common variants that influence neurodevelopmental traits in the general population. This suggests that fully understanding the genetic architecture of these disorders will require considering the full spectrum of alleles from those unique to an individual to those shared across continents.

## Methods

### DDD cohort phenotypes

Recruitment and phenotyping of DDD patients is described in detail elsewhere ^6,7^. The DDD study has UK Research Ethics Committee approval (10/H0305/83, granted by the Cambridge South Research Ethics Committee and GEN/284/12, granted by the Republic of Ireland Research Ethics Committee). Families gave informed consent for participation. Briefly, the DDD study recruited patients with a previously undiagnosed developmental disorder, in the UK and Ireland. Patient phenotypes were systematically recorded by clinical geneticists using Human Phenotype Ontology (HPO) terms in a central database, DECIPHER ^22^.

The DDD cohort is very heterogeneous in terms of patient phenotypes, and so we narrowed our analyses to singleton patients and trios where the proband had at least one of the following HPO terms or daughter terms of: abnormal metabolic brain imaging by MRS (HP:0012705), abnormal brain positron emission tomography (HP:0012657), abnormal synaptic transmission (HP:0012535), abnormal nervous system electrophysiology (HP:0001311), behavioural abnormality (HP:0000708), seizures (HP:0001250), encephalopathy (HP:001298), abnormality of higher mental function (HP:0011446), neurodevelopmental abnormality (HP:0012759), abnormality of the nervous system morphology (HP:0012639). This “neurodevelopmental” subset included both individuals who have since recruitment to the DDD study been found to carry diagnostic exome mutations in protein-coding genes ^6,20,21,28^, and individuals who are awaiting diagnosis. We therefore define our main phenotype, “neurodevelopmental disorder risk”, as the risk of having a previously undiagnosed developmental disorder and being included in the DDD study, and having at least one neurodevelopmental HPO. In addition to HPOs, some DDD patients also had a clinical record of growth measurements such as height, birth weight and head circumference.

We counted the proportion of DDD patients with particular medically relevant HPOs, displayed in Figure 2a. Individuals with the HPO were counted using a word search of the particular HPO and it’s daughter nodes. When counting the number of distinct organ systems affected in each DDD patient (Figure 2b), we faced the issue that some HPOs fell under multiple organ systems, as for example, microcephaly which is a common term in the cohort falls under three categories: “nervous system”, “head or neck” and “skeletal system”. In order to assign each HPO into only one organ system, we first ranked organ systems based on the number of raw counts of individuals with at least one term under that system (Extended Data Table 1) in the full DDD cohort. We then looked for individuals with at least one HPO under the organ system ranked most commonly affected, and assigned these individuals an organ system count of one. We then removed these HPOs from the patients’ lists, before continuing to identify individuals with at least one HPO in the organ system ranked second most prevalently affected. We continued to count organs and remove HPOs until we had assigned all individuals a count of organs systems affected out of 19 non-overlapping systems.

### Australian DD cohort phenotypes

We obtained a replication cohort of 1,270 developmental disorder cases from South Australia, originally genotyped (using the Illumina Infinium CytoSNP-850k BeadChip) as part of routine clinical care to ascertain pathogenic copy number variants. The majority (>95%) were under 18 years old. 50-60% were recruited through clinical genetics units, and the rest through neurologists, neonatologists, paediatricians and cardiologists. Based on reviewing information on the request forms, the majority of patients had developmental delay/intellectual disability and malformations involving at least one organ (e.g. brain, heart, and kidney). 15-20% were recruited as neonates with multiple malformations involving brain, heart and/or other organs, and were too young to be diagnosed with developmental delay/intellectual disability.

### Datasets and Quality Control

We genotyped 11,304 patients and 930 full trios recruited to the DDD study, on Illumina HumanCoreExome and HumanOmniExpress chips, respectively. Genotyping was carried out by the Wellcome Trust Sanger Institute genotyping facility. As controls for the discovery GWAS, we used genotype data for 10,484 individuals from the UK-based Understanding Society (UKHLS)^29,30^. Recruitment to this study was carried out through UK-wide household longitudinal survey. For replication, we obtained GWAS data from a cohort of developmental disorder cases from South Australia and population-matched controls from the Brisbane Longitudinal Twin Study (Queensland Institute of Medical Research ^31,32^). All data were on GRCh37, and detailed information of genotyping chips is shown in Supplementary Table 1.

We performed variant and sample quality control for each dataset separately. Briefly, we removed variants and samples with high data missingness, samples with high or low heterozygosity to control for admixture and inbreeding, and removed sample duplicates (steps described in detail in Supplementary Table 2). We defined sample ancestry based on a projection principal component (PCA) analysis using PLINK with 1000 Genomes Phase 3 populations, using variants with a minor allele frequency (MAF) of ≥10%. For the HumanCoreExome data and the Australian data, we removed rare variants MAF≤0.5% before imputation. For analyses described in this paper, we carried forward individuals of European ancestry, defined by selecting samples clustering around the 1000 Genomes Great British (GBR) samples in the PCA (Extended Data Figure 1). We then removed one individual from pairs of related individuals (alleles identical by descent >12%, using PLINK) from the case-control cohorts. Individuals in the discovery cohort were not related to the independent DDD trios.

### Phasing and imputation

After sample and variant quality control, we imputed all datasets in order to boost the coverage of the genome for association testing and to increase overlap of datasets genotyped on different chips. The discovery GWAS cohorts genotyped on the HumanCoreExome backbone were phased and imputed together using variants that intersected between the different versions of the chip. Trios were phased and imputed in a second batch, due to the small number of overlapping variants between the HumanOmniExpress and the HumanCoreExome chips. We phased and imputed the Australian GWAS data in a third batch, using variants that intersected between the CytoSNP-850K chip and the Illumina 610K chip. We used the Sanger Institute Imputation Service ^33^ to carry out phasing and imputation, using Eagle2 (v2.0.5) ^34^ and PBWT ^35^ respectively, selecting the Haplotype Reference Consortium as the reference panel (release 1.1, chr1-22, X) ^33^.

### Discovery GWAS of developmental disorder risk

We carried out genome-wide association study for developmental disorder risk in the discovery neurodevelopmental set of 6,987 cases and 9,270 controls of European ancestry-only, using BOLT linear mixed models ^36^ with sex as a covariate. We included in our analysis genotyped variants or high-confidence imputed variants (INFO≥0.9) with a MAF of ≥5%.

### SNP heritability

From the discovery GWAS summary statistics, we removed the MHC region (chromosome 6 region 26-34MB), and estimated trait heritability using LDSC ^8^ in LD Hub ^37^. Given the ascertainment of the DDD neurodevelopmental cases in this study, estimating the true population prevalence was not feasible. We therefore estimated single nucleotide polymorphism (SNP) heritability for our discovery GWAS on the liability scale for a range of prevalences between 0.2% and 2%, and found that SNP heritability varies from 5.5% (SE=1.5%) to 9.1% (SE=2.5%). We report heritability assuming a prevalence of 1% in the population. Heritability on the observed scale in our discovery GWAS was 13.8% (SE=3.7%).

### pTDT

We used the pTDT method, described in ^11^, to investigate transmission disequilibrium of effect alleles for traits within DDD trios, using imputed genotype data. Briefly, the test compares the means of two polygenic score distributions: one comprising of scores of the probands, and the other of the average parent-pair scores. The test is equivalent to a one-sample t-test, assessing whether the mean of score distribution in probands deviates from the mean of parent-pair score average. We report a one-sided p-value for over-transmission.

### Genetic correlation

We carried out genetic correlation of the developmental disorder risk discovery GWAS against multiple published traits using bivariate LDSC ^15^. For traits included in LD Hub we used the online server, and for traits not included in LD Hub we used the LDSC software. For genetic correlation with developmental disorder risk, we pre-selected a range of different types of traits and diseases: traits relating to cognitive performance, education, psychiatric traits and diseases, anthropometric traits and non-brain related traits and diseases. Ninety-five percent confidence intervals in Figure 3 are shown before correction for multiple testing. We set the significance threshold to p<0.0026 (0.05/19 tests).

### Partitioned heritability

We used partitioned LDSC ^18^ to look for enrichment of heritability in cell type groups and functional genomic categories. To do this we used the baseline model LD scores and regression weights available online. For cell type groups and functional categories we set the significance threshold to P<0.005 (0.05/10 tests) and P<9.6x10^−4^ (0.05/52 tests), respectively.

### Polygenic scores

We constructed polygenic scores using summary statistics from our developmental disorder risk GWAS and seven published GWAS (educational attainment ^16^, intelligence ^17^, schizophrenia ^38^, autism ^9^, intracranial volume ^39^, height ^40^ and birth weight ^41^). For all traits, we included only variants that had a MAF≥5% and were directly genotyped or imputed with high confidence (INFO≥0.9) in the respective study cohort (discovery case-control, trios or Australians). To construct the polygenic scores for individuals, we then multiplied the variant effects (betas) with the individual’s allele counts. For imputed variants, we used genotype probabilities rather than hard-called allele counts. To find independent variants for our scores, we pruned variants intersecting the original study summary statistics and our GWAS data using PLINK, by taking the top variant and removing variants within 500kb and that have r^2^≥0.1 with the top variant. We then repeated the process until no variant had a P-value below a pre-defined threshold, which we based on prior knowledge of variance in the phenotype explained. For developmental disorder risk score, we chose a P-value threshold which resulted in a score that was most strongly associated with case/control status in an independent subset of DDD patients. Specifically, we repeated our developmental disorder risk GWAS having removed a random subset of 20% of cases and controls, then calculated a score in this leave-out subset, and performed a logistic regression to assess association of case-control status with the score. The threshold P<1 performed best in two independent permutations. When deciding the P-value thresholds for published GWAS, we used the threshold that had been found to explain the most variation in other published studies for the trait (years in education P<1 ^14^, intelligence ^17^, schizophrenia P<0.05 ^38^, autism P<0.1 ^11^). For traits which we had phenotype data for in the DDD, we used thresholds that explained the most variation in DDD cases (intracranial volume P<1, birth weight P<0.01, height P<0.005). Thresholds and the number of variants used for each score are shown in Extended Data Tables 4-6. All scores were normalised to a mean of 0 and variance of 1. To test for association between trait and score, we used R (version 1.90b3) to perform logistic regression for binary traits and linear regression for quantitative traits, including the first ten principal components from the ancestry PCA to control for possible population stratification.

In order to assess power for detecting differences in scores between diagnosed and undiagnosed patients, we tested the hypothesis that diagnosed patients were effectively a random sample of controls with respect to their polygenic scores. Specifically, we randomly sampled 1,127 controls (i.e. the same number as we had diagnosed patients) and compared the polygenic scores between them and the undiagnosed patients using logistic regression. We repeated this 10,000 times and determined the proportion of times we detected a significant difference P<0.007 (P<0.05/7 correcting for seven polygenic scores) as proxy for power. For educational attainment, this was 99.1% of simulations, 93.6% for schizophrenia, and 61.2% for intelligence.

The schizophrenia PGC-CLOZUK study included some controls from the Australian cohort used in our study, and therefore we ran polygenic score analyses in the Australians using summary statistics from PGC-CLOZUK (obtained through personal communication from A. Pardinas) after these samples had been removed.

### Subsetting the DDD cohort

We defined a set of patients with an exonic diagnosis and a set with no likely diagnostic variants. This was based on the clinical filtering procedure described in ^6^, which focuses on identifying rare, damaging variants in a set of genes known to cause developmental disorders (https://www.ebi.ac.uk/gene2phenotype/), that fit an appropriate inheritance mode. Variants that pass clinical filtering are uploaded to DECIPHER, where the patients’ clinicians classify them as “definitely pathogenic”, “likely pathogenic”, “uncertain”, “likely benign” or “benign”. This process of clinical classification is necessarily dynamic as new disorders are identified and patients manifest new phenotypes. Our “diagnosed” set consists of 1,127 patients who fulfilled one of these criteria: a) amongst the diagnosed set in a recent reanalysis of the first 1,133 trios ^42^, or b) had at least one variant (or pair of compound heterozygous variants) rated as “definitely pathogenic” or “likely pathogenic” by a clinician, or c) had at least one variant (or pair of compound heterozygous variants) in a class with a high positive predictive value that passed clinical filtering but had not yet been rated by clinicians. We considered *de novo* or compound heterozygous loss-of-function (LoF) variants to have high positive predictive value, since of the ones that had been rated clinicians, 100% of compound heterozygous LoFs and 94.% of *de novo* LoFs had been classed as “definitely” or “likely pathogenic”. Our “undiagnosed” set consists of 2,479 patients who had no variants that passed our clinical filtering, or in whom the variants that had passed clinical filtering had all been rated as “likely benign” or “benign” by clinicians, or who were amongst the “undiagnosed” set in the first 1,133 trios that have previously been extensively clinically reviewed ^6^. Note that our diagnosed versus undiagnosed analysis excludes 3,375 patients who had one or more variants that passed clinical filtering in a class with a relatively low positive predictive value, but that have not yet been rated by clinicians.

We defined patients to present with autistic behaviour if their phenotype included autistic behaviour (HP:0000729) or any of its daughter nodes. We defined patients as having “mild/moderate intellectual disability or delay” if their HPO phenotypes included borderline, mild or moderate intellectual disability (HP:0006889, HP:0001256, HP:0002342) and/or mild or moderate global developmental delay (HP:0011342, HP:0011343). Patients were included in the “severe ID or delay” set if they had severe or profound intellectual disability (HP:0010864, HP:0002187) and/or severe or profound global developmental delay (HP:0011344, HP:0012736). We excluded patients with ID or global developmental delay of undefined severity.

## Extended Data Figures

**Extended Data Figure 1.**
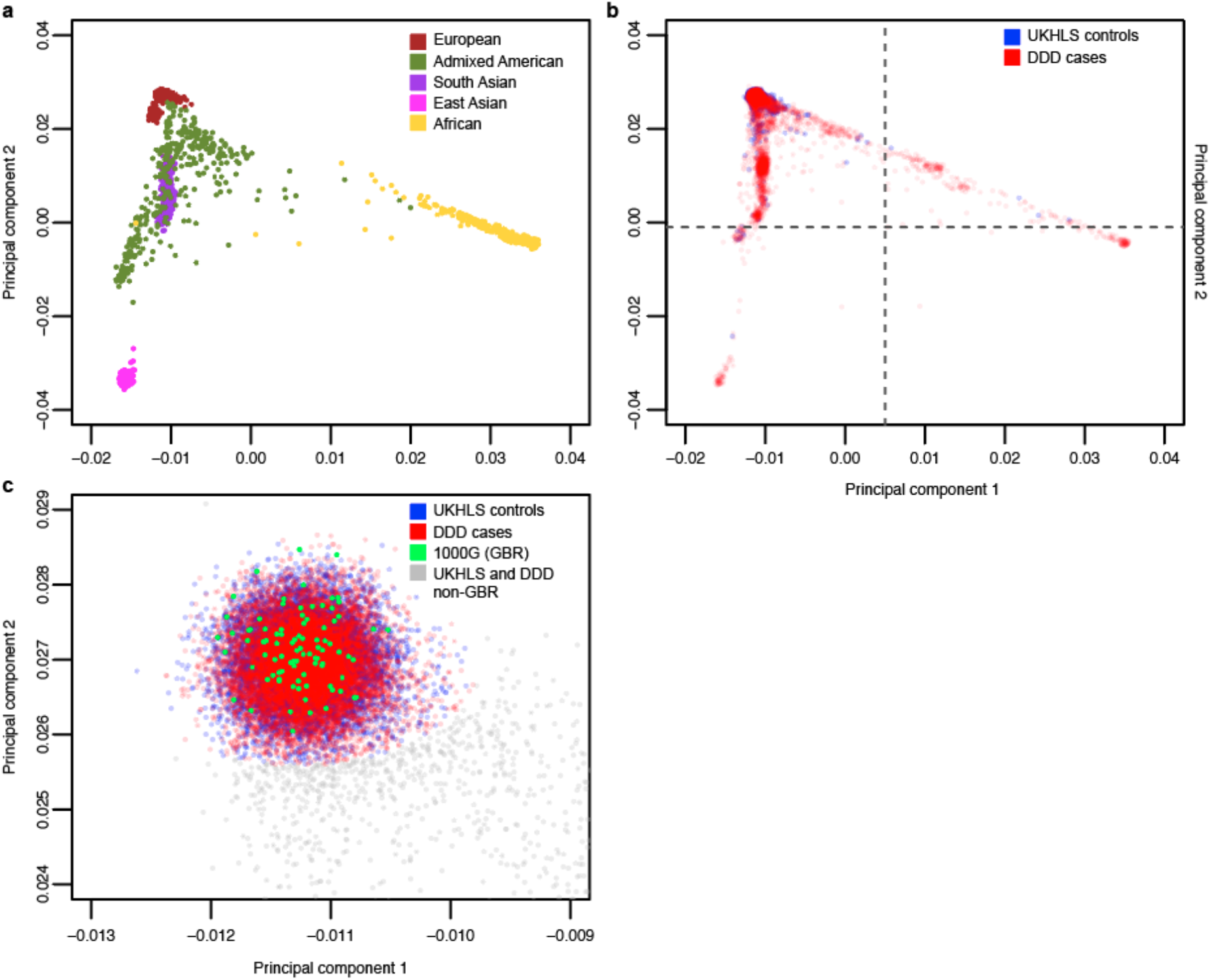
Ancestry principal components analysis of discovery GWAS cases and controls. a. Reference samples from 1000 Genomes Phase 3, coloured by the five super populations. b. Principal component projection of DDD patients, parents and UKHLS controls on 1000 Genomes samples. All DDD individuals with genotype data are plotted on top of UKHLS controls. Dotted lines show the cutoffs used to exclude East Asian and African ancestry samples before imputation. Samples in the upper right-hand square were carried forward for imputation. c. DDD patients and UKHLS controls that were selected for analyses based on proximity to 1000 Genomes GBR samples. Grey samples were excluded from further analyses.

**Extended Data Figure 2.**
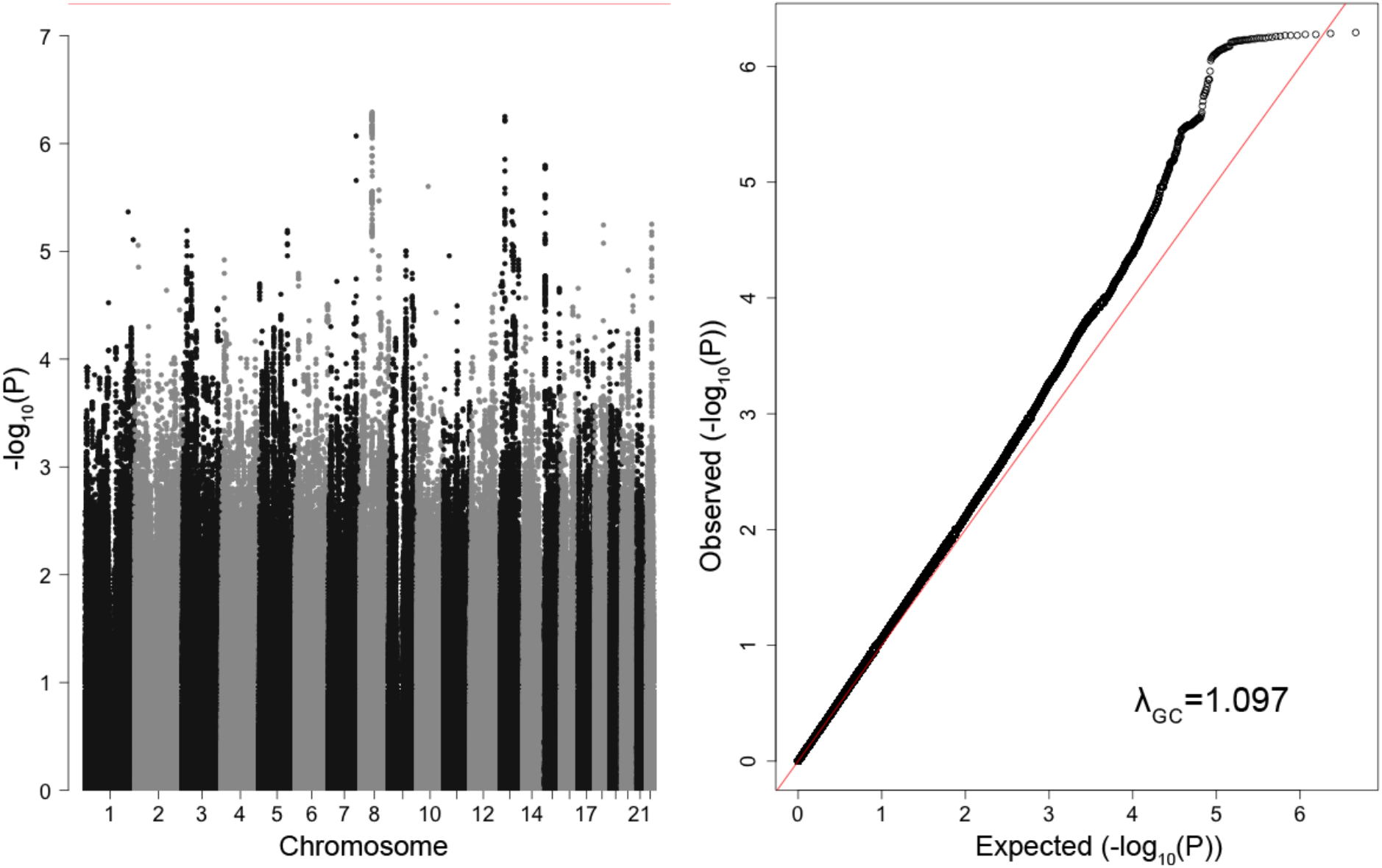
Discovery GWAS of developmental disorder risk. a. Manhattan plot of developmental disorder discovery GWAS, with 6,987 DDD cases (GBR ancestry) and 9,270 ancestry-matched UKHLS controls, including 4,733,931 variants MAF≥5%. Red line = threshold for genome-wide significance (P=5x10^−8^). b. Quantile-quantile plot of developmental disorder discovery GWAS. Red line = expected values under the null.

## Extended Data Tables

**Extended Data Table 1. Proportions of DD patients who have at least one HPO term belonging to a particular organ system category**.

**Extended Data Table 2. Genetic correlations between neurodevelopmental disorder risk and a range of traits**.

**Extended Data Table 3. Enrichment of developmental disorder risk heritability in different cell type groups**.

**Extended Data Table 4. Polygenic score analyses comparing 1,266 Australian intellectual disability cases and 1,688 controls**.

**Extended Data Table 5. Polygenic score analyses comparing DDD patients with an exome diagnosis (N=1,127) against undiagnosed patients (N=2,479)**.

**Extended Data Table 6. Polygenic score analyses in the DDD study**.

## Supplementary Tables

**Supplementary Table 1. Summary information for samples and variants genotyped on different DNA chips**.

**Supplementary Table 2. Summary of sample and variant quality control parameters used**.

**Supplementary Table 3. Enrichment of developmental disorder risk heritability in overlapping functional categories**.

## Acknowledgements

We thank the families for their participation and patience; the DDD study clinicians, research nurses and clinical scientists in the recruiting centres for their hard work and perseverance on behalf of families. The DDD study presents independent research commissioned by the Health Innovation Challenge Fund [grant number HICF-1009-003], a parallel funding partnership between Wellcome and the Department of Health, and the Wellcome Sanger Institute [grant number WT098051]. The views expressed in this publication are those of the author(s) and not necessarily those of Wellcome or the Department of Health. The study has UK Research Ethics Committee approval (10/H0305/83, granted by the Cambridge South REC, and GEN/284/12 granted by the Republic of Ireland REC). The research team acknowledges the support of the National Institute for Health Research, through the Comprehensive Clinical Research Network. This study makes use of data generated by the DECIPHER community. A full list of centres who contributed to the generation of the data is available from http://decipher.sanger.ac.uk and via email from. Funding for the project was provided by the Wellcome Trust.

We used data from Understanding Society: The UK Household Longitudinal Study, which is led by the Institute for Social and Economic Research at the University of Essex and funded by the Economic and Social Research Council (Grant Number: ES/M008592/1). The data were collected by NatCen and the genome wide scan data were analysed by the Wellcome Trust Sanger Institute. Information on how to access the data can be found on the Understanding Society website https://www.understandingsociety.ac.uk/. Data governance was provided by the METADAC data access committee, funded by ESRC, Wellcome, and MRC. (Grant Number: MR/N01104X/1).

Australian controls from the Brisbane Longitudinal Twin Study were collected and genotyped with grants from the National Health and Medical Research Council.

We thank Antonio Pardiñas for producing the PGC-CLOZUK summary statistics without the Australian controls.

## Author contributions

Study design: J.C.B., C.F.W., D.R.F., H.V.F. and M.E.H.

Data analysis and methods: M.E.K.N., H.C.M., D.L.R., G.G., M.K., J.M., and E.J.R.

Australian data collection: S.Y., J.G. and N.G.M.; Australian data preparation: K.M. and S.G. Wrote paper: M.E.K.N., H.C.M. and J.C.B.

Analytical supervision: J.C.B., H.C.M.

Project supervision: J.C.B.

## Data availability

The datasets generated during and/or analysed during the current study will be available through EGA.

